# Thinking past owl vs owl: The barred owl invasion threatens ecological communities in the western United States

**DOI:** 10.1101/2025.10.31.685899

**Authors:** Daniela Arenas-Viveros, Emma Fehlker Campbell, Amy L. Munes, Hollis Howe, Hermary M. Gonzales, J. Mark Higley, Daniel F. Hofstadter, Brendan K. Hobart, Greta M. Wengert, Angela Rex, Brian P. Dotters, Kevin N. Roberts, Christina P. Varian, M. Zachariah Peery, Emily D. Fountain

**Affiliations:** Department of Forest and Wildlife Ecology, University of Wisconsin – Madison, Madison, Wisconsin, USA; Hoopa Tribal Forestry, Hoopa, California, USA; Intergral Ecology Research Center, Blue Lake, California, USA; Sierra Pacific Industries, Anderson, California, USA

**Keywords:** Invasive species, ecosystemic impacts, Pacific Northwest, threatened species, amphibian conservation, *Strix varia*

## Abstract

Anthropogenic land use and climate change are facilitating intracontinental species range expansions, leading to novel ecological communities and interactions. Although invasive range-expanders threaten native species, lethal management can be challenged based on the notion that range-expanders are native and therefore less likely to precipitate ecological damage. The anthropogenically-mediated range expansion of barred owls (*Strix varia*) from eastern to western North America exemplifies this controversy: a recent U.S. Fish and Wildlife Service plan to cull barred owls for the benefit of spotted owls (*Strix occidentalis*) ignited public outcry and resistance. Here, we assessed barred owls’ potential invasiveness and broader effects on biodiversity by using molecular methods to characterize the diet of 788 individuals in California, Oregon, and Washington. Barred owls consumed 162 vertebrate and invertebrate prey species, including 29 species with federal- or state-level conservation status. Our findings suggest that barred owls may (1) threaten smaller native predators via intraguild predation and competition for shared prey; and (2) further imperil amphibian communities already threatened by a suite of environmental stressors. We thus contend that barred owls function as an invasive species in their expanded range. The Precautionary Principle suggests that large-scale lethal management is warranted to curb barred owls’ impacts on not just spotted owls but broader ecological communities in the forests of the Pacific Northwest. More broadly, our study highlights the significant biodiversity loss that could result from range expansions of a generalist predator and provides a roadmap for quantifying putative impacts.

## 1 | INTRODUCTION

In recent centuries, invasive species have been a leading cause of species extinctions and biodiversity loss globally (Mollot et al., 2017; Linders et al., 2019). Traditionally, invasive species have been defined as non-native species transported by humans to a new ecosystem where they cause negative ecological, environmental, or economic impacts (Blackburn et al., 2011; Wallingford et al., 2020). More recently however, anthropogenic land use and climate change have been transforming landscapes and, as a result, species dispersal, range shifts, and range expansions into new environments are becoming more common (Montràs-Janer et al., 2024; Wallingford et al., 2020; Weiskopf et al., 2020). Species extending their distribution by taking advantage of human-mediated changes, hereafter “range-expanders” (Brodie et al., 2025), may be as detrimental to ecosystems as species introduced by human transport. However, they are often overlooked due to public perception and focus on geographic origin over impacts (Davis et al., 2011; Warren, 2023). Nevertheless, range-expanders have the potential to precipitate functional and structural changes in communities that result in biodiversity declines (e.g. nine-banded armadillos (*Dasypus novemcinctus*) (Larreur et al., 2025), mountain pine beetle (*Dendroctonus ponderosae*) (Creeden et al., 2014), and striped field mouse (*Apodemus agrarius*) (Tulis et al., 2023)). With the rapid pace of climate and land-use change, the frequency of human-facilitated range expansions will inevitably increase—as will the need for difficult decisions about when, where, and how to manage the potential detrimental ecological effects of range-expanding species.

The distinction between invasive species transported by human agency and range-expanding species aided by anthropogenic disturbances can lead to apprehension regarding whether human intervention is necessary (Essl et al., 2019; Davis et al., 2011; Pecl et al., 2017). This apprehension can create resistance to the management of species the public perceives to be expanding their range “naturally”, even when the range expansion was facilitated by human activities (e.g., Canada geese (*Branta canadensis*) in the United States and European hedgehogs (*Erinaceus europaeus*) in the Scottish islands) (Crowley et al., 2017; Sharp et al., 2011). Thus, whether a range-expanding species should be classified as invasive (Nackley et al., 2017; Tong et al., 2018) and the extent to which management intervention is warranted, ultimately depends on the nature of its ecological impacts to recipient communities in conjunction with assessments of feasibility (Essl et al., 2019). One way to facilitate decision-making is to assess and communicate the effects of range-expanding species when they become invasive, including but not limited to their effects on native species of conservation concern (Pecl et al., 2017).

The anthropogenically-mediated expansion of barred owls (*Strix varia*) from eastern to western North America exemplifies the debate surrounding invasive versus range-expanding species because human-induced changes to the landscape (e.g., woody encroachment of prairies) enabled this charismatic raptor to spread to biogeographically disparate regions (Livezey, 2009). The arrival of barred owls to the Pacific Northwest, where they have now reached high densities, has the potential to trigger cascading effects across trophic levels (Holm et al., 2016). To date, the most examined consequence of the arrival of barred owls is their competition with northern and California spotted owls (*Strix occidentalis caurina and S. o. occidentalis*, respectively) (Wood et al., 2020; Wood et al., 2021; Wiens et al., 2014; Yackulic et al., 2019; Dugger et al., 2016), two subspecies that barred owls will render extinct without intervention (USFWS 2024). However, there is also cause to expect broader ecological effects through novel pressures on prey, competition with native predators, and consequent effects on ecosystem functions (Holm et al., 2016). Novel top-down pressure on prey could lead to population declines, extirpations, and extinctions of species, particularly those already threatened by other environmental stressors (Haines et al., 2024; Triay-Portella et al., 2022). Furthermore, predators such as fisher (*Pekania pennanti*) and goshawk (*Accipiter gentilis*) may encounter direct competition for prey resources, while others such as western screech owls (*Megascops kennicottii*), northern harrier (*Circus hudsonius*), ringtail (*Bassariscus astutus*), and spotted skunks *(Spilogale gracilis*) may face intraguild predation by barred owls (Rugg et al., 2023; Tosa et al., 2022; Livezey, 2007; Acker, 2012).

Recently, to mitigate the impacts of barred owls on spotted owls and potentially other listed species, the U.S. Fish and Wildlife Service (USFWS) developed a strategy involving the lethal removal of large numbers of barred owls in California, Oregon, and Washington (USFWS 2024). This proposal ignited an outcry and resistance from animal rights groups and the public (Animal Wellness Action, 2024). Alongside moral and feasibility objections, opponents assert that barred owls are native range-expanders, not an invasive species, and therefore pose no threat to western forest ecosystems. Despite several studies demonstrating that lethal removals can curb barred owl densities and promote spotted owl populations (Hofstadter et al., 2022; Diller et al., 2016; Wiens et al., 2021; Hobart et al., 2025), as well as conservation groups and scientists explaining the merits and feasibility of the strategy (The Associated Press, 2024; Birds Connect Seattle, 2024; Dumbacher and Franklin, 2025), the controversy persists (see selected press coverage in the supporting information from Hobart et al. (2025)). Notably, most criticisms and coverage of the barred owl management strategy focus largely or solely on competition between barred and spotted owls, with little attention given to the broader consequences barred owls may have on native species and their ecological communities.

Previous dietary studies have shown that barred owls are generalist predators, consuming a wide range of vertebrates and invertebrates during a foraging cycle (Wiens et al., 2014; Baumbusch, 2023; Kryshak et al., 2022; Livezey, 2007). Despite the large number of prey species already identified, the visual techniques used in most of these studies are biased towards prey items that leave indigestible remains like bones and exoskeletons (Hoenig et al., 2022; Nielsen et al., 2018; Symondson, 2002). In contrast, DNA techniques offer unprecedented levels of taxonomic resolution and have greater sensitivity to detect rare species and soft or highly degraded items (Nielsen et al., 2018; Symondson, 2002).

This increased sensitivity facilitates the detection of trophic interactions that might otherwise remain undetected, including the consumption of rare species that are likely to constitute a small proportion of a predator’s diet (e.g., threatened and endangered species; (Tosa et al., 2022)). Here, we characterize barred owl prey consumption using molecular methods, leveraging the largest number of individuals sampled to date across their invasive range from Washington and Oregon to the leading edge in California.

Specifically, we provide a detailed list of prey items consumed with special emphasis on native species of conservation concern. Identifying the extent to which barred owls consume already at-risk species in the western United States is key for guiding policy in this high-profile conservation conflict.

## 2 METHODS

### 2.1 Study Area and Barred Owl Removals

We assessed barred owl diet using DNA sequencing methods applied to the intestinal contents of lethally removed individuals (n = 788) collected year-round across a wide range of different forest types and conditions in Washington, Oregon, and California from 2015 to 2024. We removed barred owls using well established methods (Diller et al., 2016) demonstrated to be humane for barred owls (Dumbacher and Franklin, 2025). In California, our sampling encompassed the invasion front, extending from Sonoma and Marin Counties in the west to Placer and El Dorado Counties in the east, and northward to Del Norte County. Samples were also collected in Klamath County and along the coastal region in Oregon, and in the more xeric forests of Kittitas County, Washington (Figure 1).

**Figure 1.**
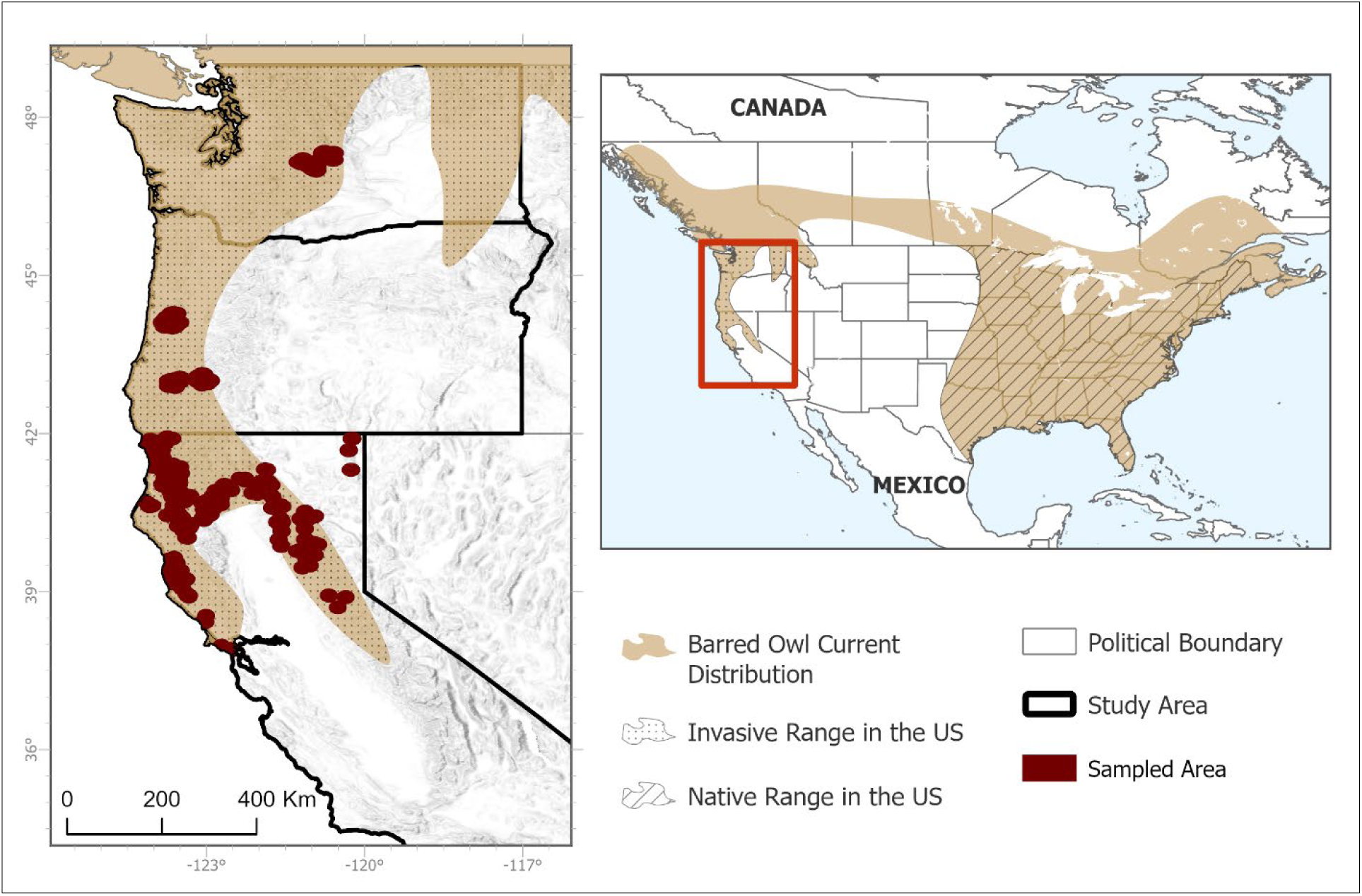
Current distribution of the barred owl (*Strix varia*) in North America, overlaid with locations where individuals were sampled. In the left panel, our study area includes their invasive range in the United States in California, Oregon and western Washington.

All removals were conducted under appropriate permits. For the University of Wisconsin: Federal Migratory Bird Permit MB24592D, California Department of Fish and Wildlife Scientific Collection Permit S-193180001-19337, and UW IACUC protocol AA006106. For the Hoopa Valley Reservation: Federal Migratory Bird Permits MB14305B-0, MB14305B-2, MB14305B-3, MB14305B-6, and MBPER0021702. For SPI: Federal Migratory Bird Permit MB53229B and California Department of Fish and Wildlife Scientific Collection Permit S-213440002-21347-001. In Oregon and Washington: Federal Migratory Bird Permit MB14305B-0 and State Scientific Collection Permits MB14305B-5 and HENSON 18-261, respectively.

### 2.2 Genetic Methods and Analysis

To identify the prey consumed by barred owls, we followed the metabarcoding and bioinformatic methods outlined by Kryshak et al. (2022). Briefly, we extracted DNA from intestinal contents and then PCR-amplified 150–200 bp fragments of mitochondrial DNA from major animal taxonomic groups: insects, arachnids, gastropods, annelids, crustaceans, and all vertebrates. We equimolarly combined the cleaned PCR amplifications for each individual and then attached unique i5 and i7 Illumina barcodes. Subsequently, we cleaned and combined all tagged PCR products equimolarly for paired-end sequencing on an Illumina MiSeq platform at the Genomics and Cell Characterization Core Facility, University of Oregon. We used the dada2 (Callahan et al., 2016) plugin in QIIME2 (Bolyen et al., 2019) to quality filter and trim raw data and build consensus sequences. Molecular operational taxonomic units (MOTUs) were assigned using a custom classifier developed from a reference database of publicly available prey sequences (Sayers et al., 2022), and prey sequences derived in our laboratory from opportunistic sampling in the field. After assigning MOTUs, we excluded any species smaller than 1.5 cm to account for secondary or accidental predation (i.e., when a mesopredator eats a target prey and then is itself consumed by a secondary predator such as a barred owl (Nielsen et al., 2018; Symondson, 2002)). While we tried to remove invertebrate species that were likely secondary consumption, we used a conservative threshold to avoid removing species that we know from observation of stomach contents (Baumbusch, 2023; Angela Rex and Emily Fountain, personal communication) are targeted by barred owls (e.g., Jerusalem crickets, centipedes, wood scorpions, side-banded snails, and crayfish). We calculated (for the combined dataset and for each state) the proportion of prey species occurring in each taxonomic class as the number of species belonging to a given class divided by total number of species consumed. We also calculated the proportion of occurrence for each prey species as the number of barred owls detected consuming a species divided by the total number of prey detections across all barred owls using the R package vegan (Oksanen et al., 2022).

### 2.3 Prey Status

Once we consolidated a list of prey species consumed by barred owls, we reviewed the USFWS Environmental Conservation Online System (https://ecos.fws.gov) to identify which prey items were listed as endangered or threatened under the federal Endangered Species Act (ESA). Additionally, to establish species with special conservation considerations in each state, we reviewed the Special Animals List from the California Natural Diversity Database (CNDDB, 2025) and California’s State Wildlife Action Plan (SWAP), the Strategy Species list from the Oregon Conservation Strategy (https://www.oregonconservationstrategy.org), and Washington’s SWAP (https://wdfw.wa.gov). In California, this includes the categories “Species of Special Concern” and “Fully Protected” as established by the California Department of Fish and Wildlife (CDFW) and “Climate Vulnerable” species and “Bird of Conservation Concern” as listed in SWAP. For Oregon, we identified species considered Strategy Species as part of the state’s Conservation Strategy, a category assigned to Species of Greatest Conservation Need. In Washington, we identified species included as Species of Greatest Concern in Chapter 3 of the SWAP.

## 3 RESULTS

We detected 162 species consumed by barred owls in Washington, Oregon, and California (Table S1). On average we detected 2.97 species consumed per owl with a minimum of one species and a maximum of 17 species detected in a single owl. Mammals were the most frequently consumed class overall (46 species), followed by birds (24 species), amphibians (23 species) and insects (21 species). Among vertebrate prey, mammals accounted for the highest proportion of prey species (0.41), followed by birds (0.22), amphibians (0.21), and reptiles (0.15)—with fish being the least commonly consumed (0.01) (Figure 2A). This pattern was generally consistent among states, except in Washington where barred owls consumed a relatively high proportion of reptile species (0.20), and in Washington and Oregon where barred owls consumed a relatively low proportion of bird species (0.12 and 0.16, respectively; Figure 2A). Among non-insect invertebrates, centipedes (class Chilopoda) and worms (class Clitellata) made up a high proportion of the overall diet in Oregon and Washington (0.12 and 0.18, respectively; Figure 2B). Barred owls consumed crayfish (class Malacostrata) in California (0.04) and Washington (0.06), but not in Oregon. Nonetheless, estimated invertebrate richness should be interpreted with caution given the possibility of secondary or accidental consumption (see Methods).

**Figure 2.**
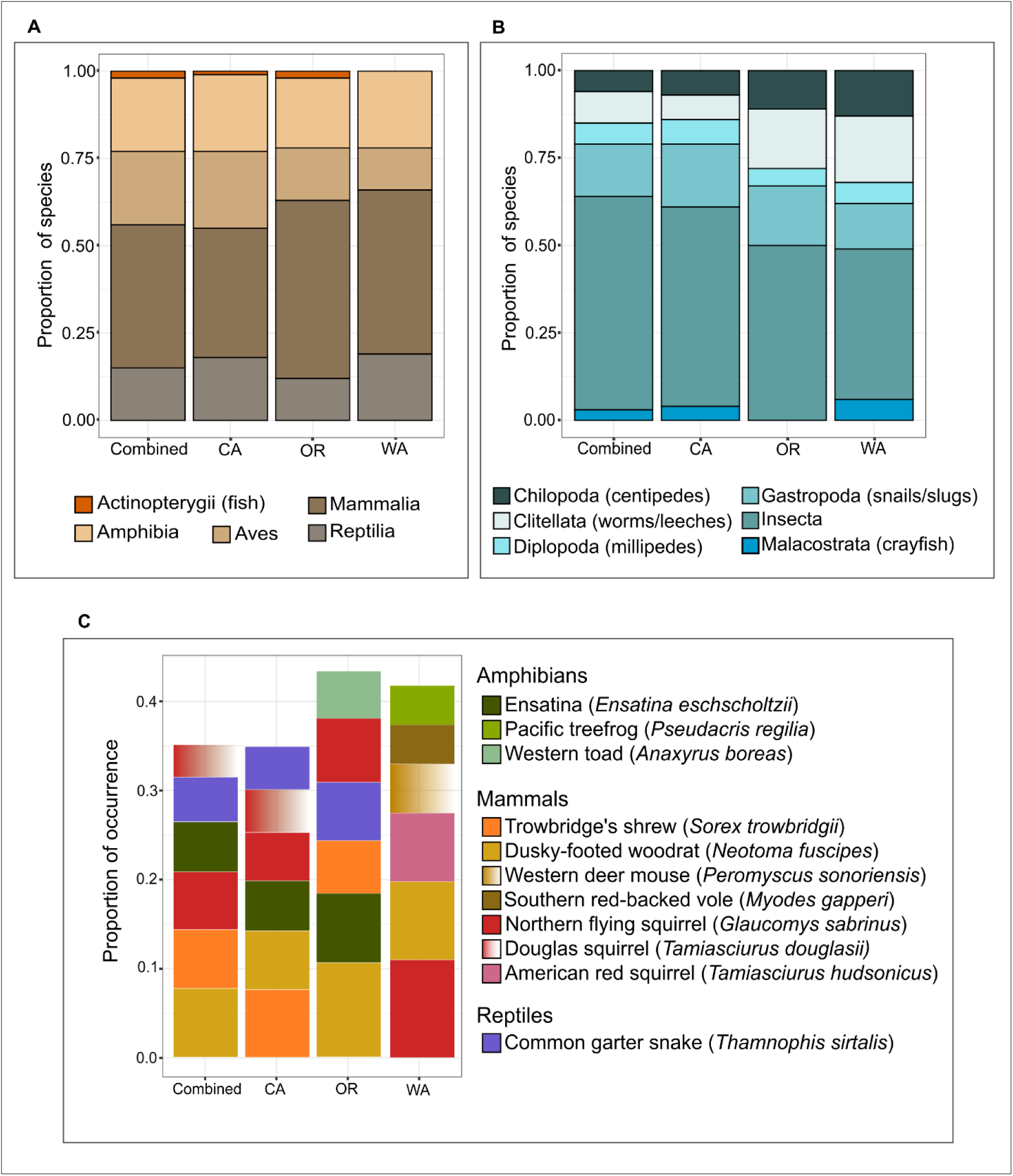
The proportion of species consumed by barred owls (*Strix varia*) in their invasive range in California, Oregon, and Washington, by class, for vertebrates (**A**) and invertebrates (**B**). Proportion of occurrence (across 788 sampled barred owls) for the six most consumed vertebrate species (**C**).

At the species level, taxa with the highest proportion of occurrence included shrews, mice, squirrels, voles, salamanders, frogs and toads, and one species of snake (Figure 2C). Dusky-footed woodrats (*Neotoma fuscipes*) and northern flying squirrels (*Glaucomys sabrinus*) were among the six most consumed species in all three states. Trowbridge’s shrew (*Sorex trowbridgii*), ensatina (*Ensatina eschscholtzii*), and common garter snake (*Thamnophis sirtalis*) were consumed commonly in California and Oregon (Figure 2C). Notably, the Western toad (*Anaxyrus boreas*) which was consumed in high proportion in Oregon, is listed by the state as a Strategy Species (see below and Table 1).

**Table 1.**
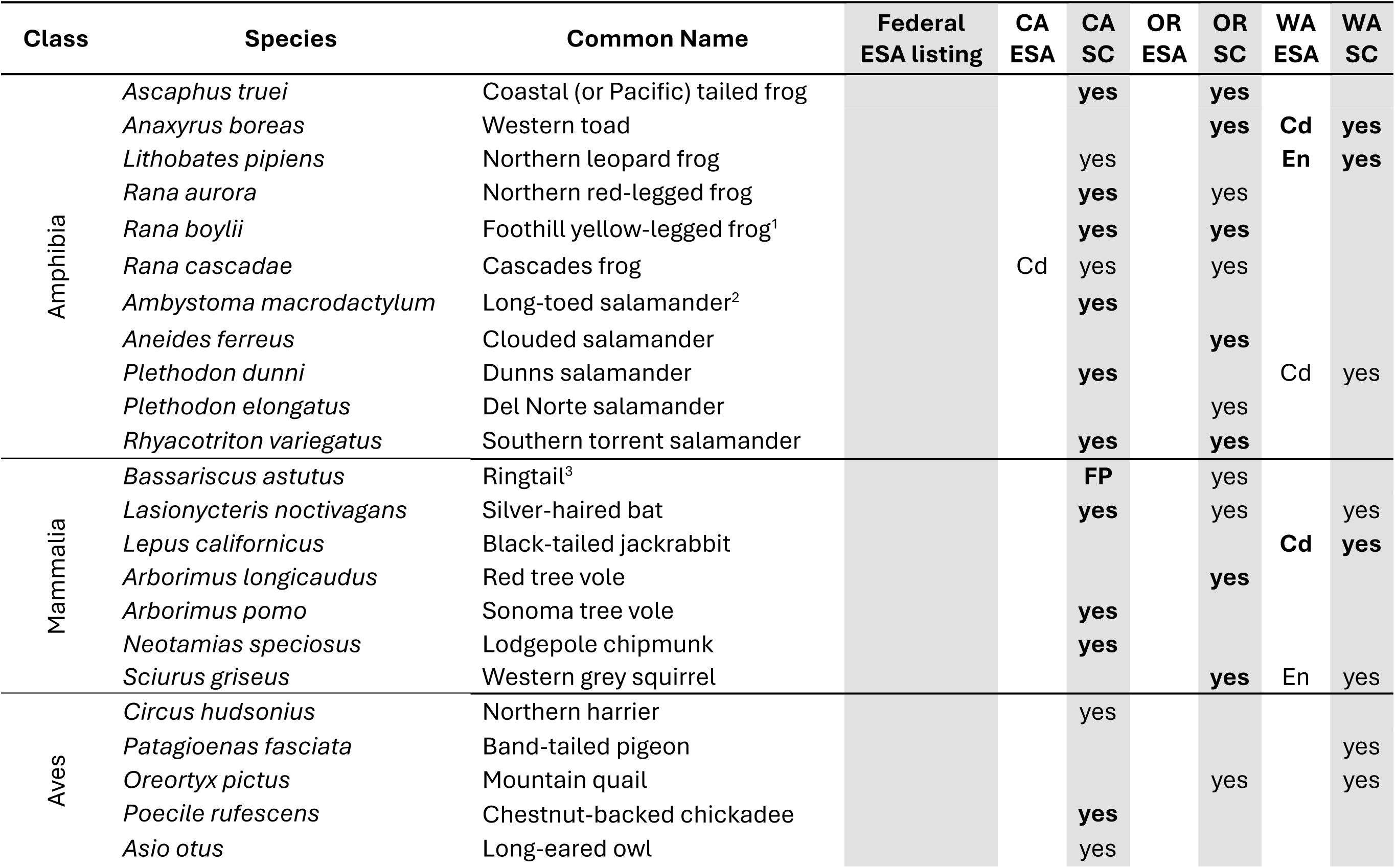
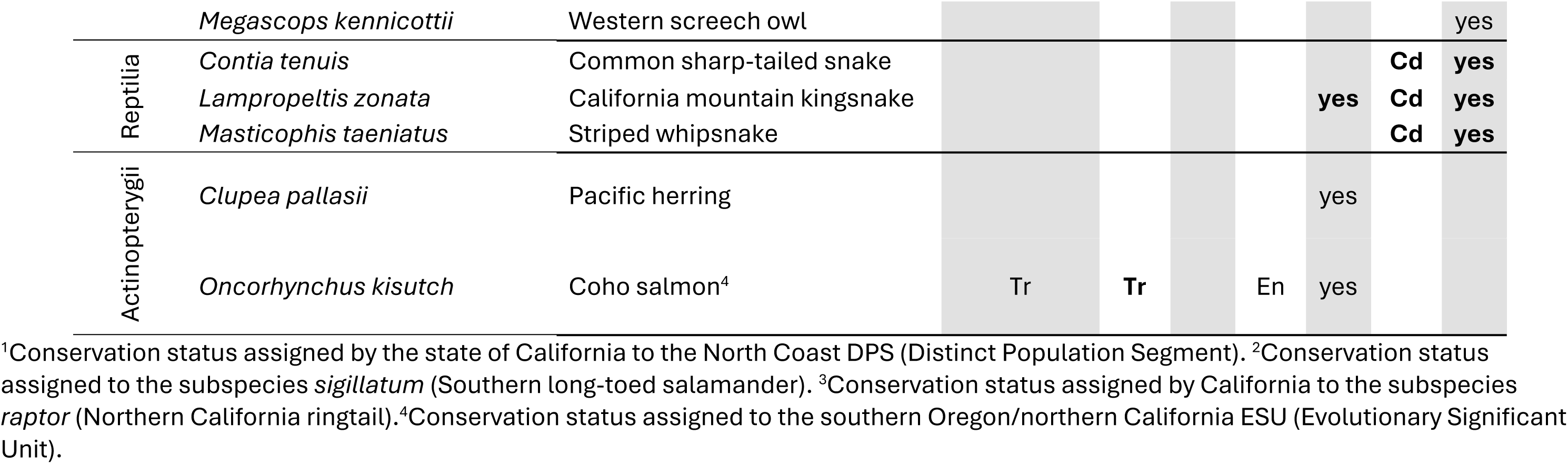
Prey species of conservation concern consumed by barred owls in their invasive range. Tr = threatened, En = endangered, Cd = candidate, FP = Fully Protected. CA = California, OR = Oregon, WA = Washington. ESA = Endangered Species Act. SC = Species of Concern (from non-ESA state-level listings). Bolded status represents instances where the prey item was consumed in the state where it is listed.

Only vertebrates had federal or state conservation status, of which 11 were amphibians, seven mammals, six birds, three reptiles, and two fish, for a total of 29 species (Table 1; Figure 3). The coho salmon (Oncorhynchus kisutch), which was detected in only a single barred owl sample, was listed as threatened at the federal ESA, and as threatened and endangered by the states of California and Oregon. The Northern leopard frog (*Lithobates pipiens*) and the Western grey squirrel (*Sciurus griseus*) were listed as endangered by the state of Washington. Species classified as candidates to be listed as endangered under a state’s ESA included the Cascades frog (*Rana cascadae*) in California, and the western toad (*Anaxyrus boreas*), Dunn’s salamander (*Plethodon dunni*), black-tailed jackrabbit (*Lepus californicus*), common sharp-tailed snake (*Contia tenuis*), California mountain kingsnake (*Lampropeltis zonata*), and striped whipsnake (*Masticophis taeniatus*) in Washington.

**Figure 3.**
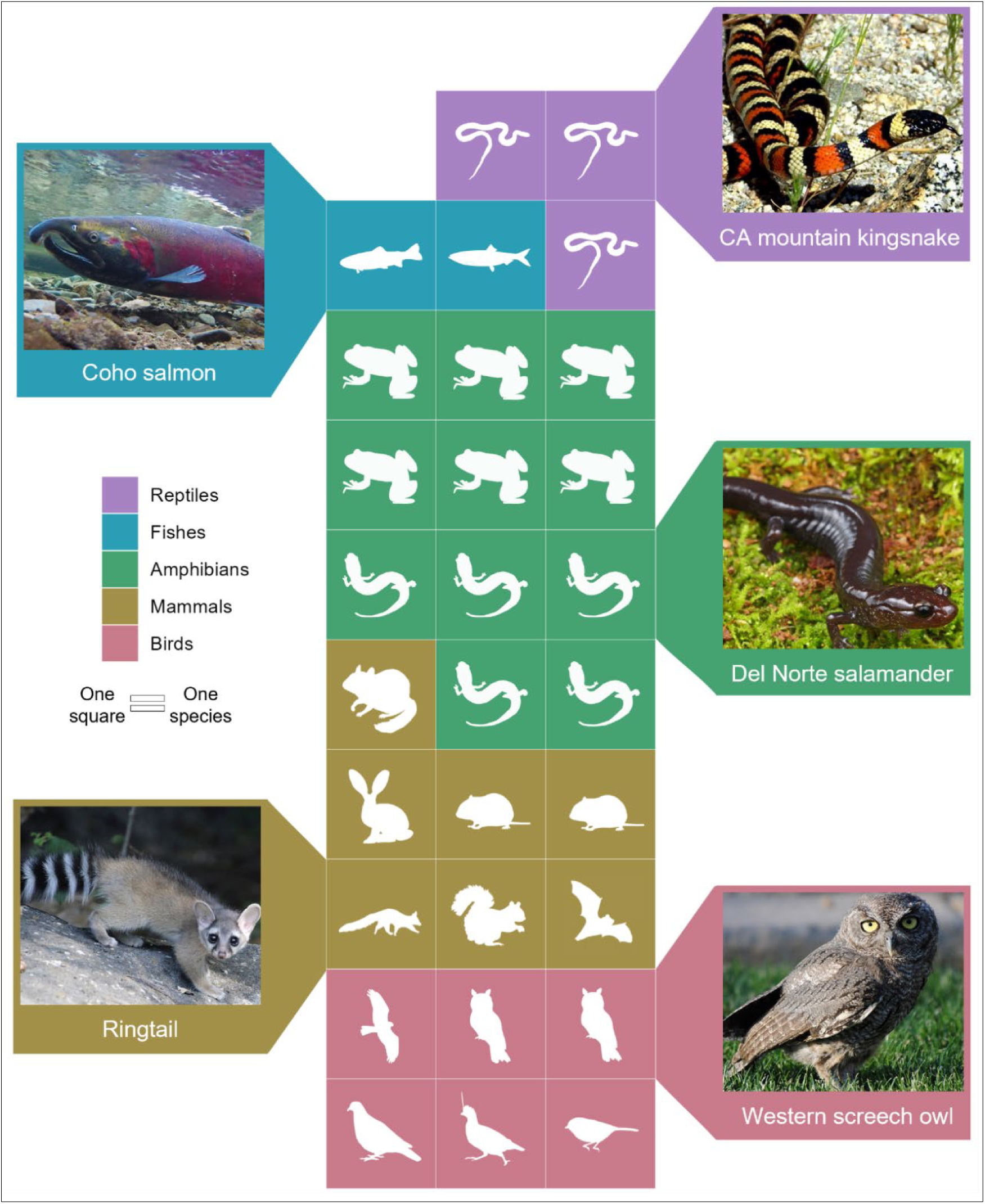
Number of species per class consumed by barred owls (*Strix varia*) in their invasive range that hold federal or state conservation concern status in the western United States. The proportion of occurrence of these species in the barred owl diet ranges from 0.001 to 0.019.

Additionally, 10 species were considered Species of Special Concern in California, four were part of the state’s SWAP, and the ringtail (*Bassariscus astutus*) was classified as Fully Protected. Sixteen species were part of the Oregon Conservation Strategy and 12 were considered Species of Greatest Concern in Washington’s SWAP (Table 1).

## 4 DISCUSSION

Our contribution represents the most taxonomically and geographically comprehensive assessment of the diet of barred owls in the western United States. The broad diet of barred owls in their invasive range (162 species spanning five vertebrate and seven invertebrate classes) indicates that this predator is likely having far-reaching impacts on native species and biodiversity in the Pacific Northwest. Top-level consumers, both native and non-native, impact biological communities through a network of direct (e.g., predation) and indirect (e.g., competition for prey resources) ecological interactions (Estes et al., 2011; Roemer et al., 2002). Barred owls are no exception, and as evidenced by our results, prey on species across trophic levels (i.e., herbivores, carnivores, omnivores), reinforcing the potential for complex ecological effects through novel and continued consumption (Holm et al., 2016). Given their high densities in the Pacific Northwest, barred owls likely exert top-down pressure on native species’ relative abundance and community structure, potentially contributing to population loss, species extinctions, and an overall erosion of biodiversity.

In our study, 29 species consumed by barred owls were of federal or state conservation concern in Washington, Oregon, and California. Invasive predators, such as barred owls, are often an additional stressor for at-risk species that are already threatened by habitat loss and other environmental factors (Doherty et al., 2016). In some instances, the pressure exerted by a non-native predator is the final stressor that triggers rapid population decline and ultimately extinction (e.g., populations of the endangered northern bettong (*Bettongia tropica*) in Australia when subjected to predation by domestic cats (Whitehead et al., 2018)). Similar outcomes could unfold throughout the invasive range of barred owls, where native species are threatened by small population sizes and fragmented distributions (e.g., western grey squirrel and ringtail (Ryan and Carey, 1995; Muchlinski et al., 2009; Gundermann et al., 2023)), habitat loss or degradation (e.g., Coho salmon, mountain quail, long-eared owl, and western toad (Crawford, 2000; Denver W Holt, 1997; Forrest et al., 2017; Weitkamp et al., 2000)), and climate change (e.g., silver-haired bat, Cascades frog, and Dunn’s salamander (Garwood and Welsh, 2007; Pelletier and Carstens, 2016; Gerald Hayes and Wiles, 2013)).

The at-risk species consumed by barred owls in this study constituted a low proportion of their diets (range: 0.001 to 0.019), but even infrequent consumption of uncommon and rare species can have emergent population-level impacts. Indeed, saw-whet, screech, and flammulated owls are uncommonly detected in barred owl diets (Wiens et al., 2014; Hamer et al., 2001; Kryshak et al., 2022), yet they have been shown to decline numerically following the arrival of barred owls (Acker, 2012) or increase following the removal of barred owls (Barry et al., 2025, Wiens et al., 2025). Our molecular assessment, while limited to species identification, highlights the potential for low-frequency interactions to disproportionately affect vulnerable populations. Additionally, the diet items detected in this study were likely consumed by owls during their most recent foraging cycle and do not contain past, longer-term predation that could have detrimental population impacts. Ultimately, the extent of these impacts on each of the species we detected will likely depend, in part, on their ecological, demographic, and natural history traits. Hence, population-level assessments of the potential numerical effects of barred owl predation on at-risk species are warranted.

Barred owls in the western U.S. may pose a particularly important threat to amphibian communities given that 11 of the 29 at-risk species consumed, and three of the six most consumed species (ensatina, Pacific treefrog, and western toad), were amphibians. Amphibians are experiencing a global extinction crisis due to habitat loss and disease, as well as other contributing environmental factors (Bolochio et al., 2020; Halliday, 2007; Bucciarelli et al., 2014). In the Pacific Northwest, where amphibian diversity is second only to that of the southeastern US, populations of the northern red-legged frog, northern leopard frog, and Cascades frog are already threatened, among others, by the presence of invasive American bullfrogs and fish (Kiesecker et al., 2001; Marc P. Hayes and Jennings, 1986; Fellers and Drost, 1993). For sensitive native species like amphibians, the presence of multiple invasive predators with similar feeding preferences can have an additive detrimental impact (Jackson, 2015). Lastly, compared to other vertebrate classes, the proportion of amphibian species consumed by barred owls was very similar across Washington, Oregon, and California, emphasizing their reliance on this group. Thus, barred owls may exert geographically pervasive effects on threatened amphibian communities within their invasive range.

In natural ecosystems, predator communities (e.g., mesocarnivores and raptors) have evolved strategies to coexist despite spatial and dietary overlaps, including foraging at different times of day and partitioning preferred prey by size, season, or type (Glen and Dickman, 2008; Young et al., 2023). However, the introduction of an invasive predator, particularly a generalist, can disrupt these ecological interactions, impacting both omnivores (e.g., raccoons, ringtails, and skunks) and carnivores (e.g., mustelids, hawks, and other owl species) (Young et al., 2023). In the Pacific Northwest, barred owls have the potential to compete for prey resources with virtually every other mammalian and avian forest predator (e.g., fisher, marten, ringtail, owls, hawks, and harriers) that feed on small mammals, amphibians, reptiles, and birds (Happe et al., 2021; Wilk and Raphael, 2017; Alexander et al., 1994; Holt and Leslie, 1996; Richardson et al., 2001). In addition, the presence of some of these predators in the diet of barred owls, including species of concern such as northern harrier, ringtail, and western spotted skunk (Tosa et al., 2022), confirms the existence of intraguild predation and suggest the possibility of predator suppression by barred owls (Young et al., 2023). Even when these species are found infrequently in barred owl diets, mesopredator populations are uniquely vulnerable to increased mortality given their low population density and reproductive capacity (Tosa et al., 2022).

Although the introduction of barred owls in the western United States is associated with several potential negative outcomes—including their widespread consumption of at-risk and sensitive species as presented here—their management has faced criticism based on feasibility, ethics, and public perception. Much of the discourse opposed to lethal management is framed within an owl-versus-owl context (i.e., barred owl versus spotted owl). We provide a more encompassing perspective on this conservation issue, which requires consensus among scientists, policymakers, and the public, beginning with the classification of barred owls in the western U.S. The terminology used in invasion ecology to refer to non-native species is ambiguous and prone to subjective interpretations. Here, we adopt the terminology from Soto et al. (2024) to classify barred owls as invasive non-natives (herein abbreviated to invasive) because 1) they arrived in western North America— where they have no evolutionary history—through range expansion following anthropogenic changes in climate and landscape (Potts, 2024; Livezey, 2009), and 2) they have established, reproduced, and are continuously expanding their range southwards, causing harmful ecological and cultural impacts by inflicting novel ecological pressures on native species through competition and predation.

Similar to Wood et al. (2020), we invoke the Precautionary Principle (Kriebel et al., 2001; Ashford et al., 1998) to suggest that lethal management of barred owls is warranted to mitigate their effects on spotted owls and to prevent broader biodiversity loss within their invasive range. The Precautionary Principle posits that management intervention, such as the removal of invasive species, should be executed despite uncertainties about their numerical effects on native species and biodiversity, particularly when inaction’s cost is high. In this case, the cost of inaction includes potential impacts, through consumption and competition, to at least 29 native at-risk vertebrate species. Furthermore, it is very likely that if they continue to expand southward, barred owls will encounter and impact other endangered populations of species they commonly consume further north, such as the San Francisco subspecies of the common garter snake (*T*. *s*. *tetrataenia*). Lethal management has proven to be effective in reducing barred owl densities from local to near-state scales, even in areas where the species has reached high densities (Diller et al., 2016; Wiens et al., 2021, 2025; Hobart et al., 2025; Hofstadter et al., 2022). Consequently, the recently published USFWS barred owl management plan, primarily developed to prevent the extinction of spotted owls, is likely to benefit a broader array of native species and potentially avert large-scale loss of biodiversity in the Pacific Northwest. Indeed, the ongoing debate about the ethics and costs of implementing the barred owl strategy would benefit from broadening beyond an “owl versus owl” perspective to acknowledge barred owls’ significant effects on biodiversity within their invasive range.

## 5 CONCLUSION

The barred owl is just one of many species expanding their geographic range in association with human activities, including several eastern species that have dispersed westward across the Great Plains and established themselves in western North America (Dumbacher and Franklin, 2025). While many range-shifting species, such as those tracking their climatic niche in a warming world, are less likely to become invasive (Urban, 2020), our findings clearly highlight the potentially significant effects that some range-expanding species can have on biodiversity within the invasive portion of their range. As humans continue to alter ecosystems, range expansions are likely to become more prevalent, necessitating difficult decisions regarding when, where, and how to intervene to ensure the conservation of biodiversity. Accordingly, decision-making will benefit from including ecological communities over a single species to assess the effects that range-expanding species exert on the invaded ecosystem. For many invasive predators, diet metabarcoding constitutes a powerful tool for assessing biodiversity impacts at the community level and should be included as part of management strategies.

## CRediT AUTHORSHIP CONTRIBUTION STATEMENT

**Daniela Arenas Viveros:** Writing – review & editing, Writing – original draft, Visualization, Validation, Methodology, Investigation, Formal analysis, Data curation, Conceptualization. **Emma Fehlker Campbell:** Writing – review & editing, Investigation, Formal analysis, Data curation. **Amy L. Munes:** Writing – review & editing, Investigation. **Hollis Howe:** Writing – review & editing, Investigation. **Hermary M. Gonzales:** Writing – review & editing, Investigation, Formal analysis, Data curation. **J. Mark Higley:** Writing – review & editing, Resources, Funding acquisition. **Daniel F. Hofstadter:** Writing – review & editing, Resources. **Brendan K. Hobart:** Writing – review & editing, Writing – original draft, Visualization, Investigation. **Greta M. Wengert:** Writing – review & editing, Funding acquisition. **Angela Rex:** Writing – review & editing, Resources. **Brian P. Dotters:** Writing – review & editing, Resources, Funding acquisition. **Kevin N. Roberts:** Writing – review & editing, Resources, Funding acquisition. **Christina P. Varian:** Writing – review & editing, Resources. **M. Zachariah Peery:** Writing – review & editing, Writing – original draft, Supervision, Resources, Funding acquisition, Conceptualization. **Emily D. Fountain:** Writing – review & editing, Writing – original draft, Validation, Supervision, Resources, Methodology, Formal analysis, Data curation, Conceptualization

## Supporting information

Complete prey list

## ACKNOWLEDGEMENTS

We would like to thank all individuals affiliated with all coauthor’s organizations who contributed to this research, particularly all removers. We also thank J. David Wiens for his contribution of samples from WA and OR and to barred owl research. We also thank the Museum of Vertebrate Zoology of University of California - Berkeley, including C. Cicero, T.L.W. Barclay, S. Medina, M. Paramonova, and A. Berger, for assistance with necropsies and prey biosamples (Loans: 2019.8892, 2023.9397), California Academy of Science for assistance with necropsies and prey biosamples (Loan 1575), and the Vertebrate Museum of Cal Poly Humboldt for prey biosamples.

