## Supplementary material for "Thinking past owl vs owl: The barred owl invasion threatens ecological communities in the western United States": Complete prey list

**Table S1.** Prey items consumed by barred owls in their invasive range in the western United States.

| Class | Order | Family | Scientific Name | Common Name |
|---|---|---|---|---|
| Amphibia | Anura | Ascaphidae | *Ascaphus truei* | Coastal tailed frog |
| | Anura | Bufonidae | *Anaxyrus boreas* | Western toad |
| | Anura | Hylidae | *Pseudacris regilla* | Pacific treefrog |
| | Anura | Hylidae | *Pseudacris sierra* | Sierran treefrog |
| | Anura | Ranidae | *Lithobates catesbeianus* | American bullfrog |
| | Anura | Ranidae | *Rana aurora* | Northern red-legged frog |
| | Anura | Ranidae | *Rana boylii* | Foothill yellow-legged frog |
| | Anura | Ranidae | *Rana cascadae* | Cascades frog |
| | Anura | Ranidae | *Lithobates pipiens* | Northern leopard frog |
| | Caudata | Ambystomatidae | *Ambystoma macrodactylum* | Long-toed salamander |
| | Caudata | Ambystomatidae | *Dicamptodon ensatus* | California giant salamander |
| | Caudata | Ambystomatidae | *Dicamptodon tenebrosus* | Coastal giant salamander |
| | Caudata | Plethodontidae | *Aneides ferreus* | Clouded salamander |
| | Caudata | Plethodontidae | *Aneides flavipunctatus* | Black salamander |
| | Caudata | Plethodontidae | *Aneides lugubris* | Arboreal salamander |
| | Caudata | Plethodontidae | *Aneides vagrans* | Wandering salamander |
| | Caudata | Plethodontidae | *Batrachoseps attenuatus* | California slender salamander |
| | Caudata | Plethodontidae | *Ensatina eschscholtzii* | Ensatina |
| | Caudata | Plethodontidae | *Plethodon dunni* | Dunns salamander |
| | Caudata | Plethodontidae | *Plethodon elongatus* | Del Norte Salamander |
| | Caudata | Plethodontidae | *Plethodon vehiculum* | Western redbacked salamander |
| | Caudata | Rhyacotritonidae | *Rhyacotriton variegatus* | Southern torrent salamander |
| | Caudata | Salamandridae | *Taricha granulosa* | Rough-skinned newt |
| Aves | Accipitriformes | Accipitridae | *Buteo lagopus* | Rough-legged hawk |
| | Accipitriformes | Accipitridae | *Circus hudsonius* | Northern harrier |
| | Anseriformes | Anatidae | *Anas clypeata* | Northern shoveler |
| | Anseriformes | Anatidae | *Anas platyrhynchos* | Mallard |
| | Anseriformes | Anatidae | *Aythya fuligula* | Tufted duck |

| Class | Order | Family | Scientific Name | Common Name |
|---|---|---|---|---|
| Aves | Columbiformes | Columbidae | *Columba livia* | Rock pigeon |
| | Columbiformes | Columbidae | *Patagioenas fasciata* | Band-tailed pigeon |
| | Columbiformes | Columbidae | *Zenaida macroura* | Mourning dove |
| | Falconiformes | Falconidae | *Falco sparverius* | American kestrel |
| | Galliformes | Odontophoridae | *Oreortyx pictus* | Mountain quail |
| | Galliformes | Phasianidae | *Bonasa umbellus* | Ruffed grouse |
| | Galliformes | Phasianidae | *Gallus gallus* | Chicken |
| | Galliformes | Phasianidae | *Meleagris gallopavo* | Turkey |
| | Passeriformes | Corvidae | *Corvus brachyrhynchos* | Common crow |
| | Passeriformes | Corvidae | *Cyanocitta stelleri* | Stellars jay |
| | Passeriformes | Paridae | *Poecile atricapilla* | Black-capped chickadee |
| | Passeriformes | Paridae | *Poecile rufescens* | Chestnut-backed chickadee |
| | Passeriformes | Parulidae | *Setophaga coronata* | Yellow-rumped warbler |
| | Passeriformes | Troglodytidae | *Troglodytes aedon* | Northern house wren |
| | Piciformes | Picidae | *Dryobates pubescens* | Downy woodpecker |
| | Strigiformes | Strigidae | *Aegolius acadius* | Northern saw-whet owl |
| | Strigiformes | Strigidae | *Asio otus* | Long-eared owl |
| | Strigiformes | Strigidae | *Bubo virginianus* | Great horned owl |
| | Strigiformes | Strigidae | *Megascops kennicottii* | Western screech owl |
| Mammalia | Carnivora | Felidae | *Felis catus* | Domestic cat |
| | Carnivora | Mustelidae | *Lontra americana* | North American river otter |
| | Carnivora | Mustelidae | *Mustela richardsonii* | American ermine |
| | Carnivora | Procyonidae | *Bassariscus astutus* | Ringtail |
| | Carnivora | Procyonidae | *Procyon lotor* | Raccoon |
| | Chiroptera | Vespertilionidae | *Myotis lucifugus* | Little brown bat |
| | Chiroptera | Vespertillionidae | *Lasionycteris noctivagans* | Silver-haired bat |
| | Didelphimorphia | Didelphidae | *Didelphis virginiana* | Virginia opossum |
| | Eulipotyphla | Soricidae | *Sorex cinereus* | Common shrew |
| | Eulipotyphla | Soricidae | *Sorex sonomae* | Fog shrew |
| | Eulipotyphla | Soricidae | *Sorex trowbridgii* | Trowbridge's shrew |
| | Eulipotyphla | Soricidae | *Sorex vagrans* | Vagrant shrew |

| Class | Order | Family | Scientific Name | Common Name |
|---|---|---|---|---|
| | Eulipotyphla | Talpidae | *Scapanus latimanus* | Broad-footed mole |
| | Eulipotyphla | Talpidae | *Scapanus orarius* | Coast mole |
| | Lagomorpha | Leporidae | *Lepus californicus* | Black-tailed jackrabbit |
| | Lagomorpha | Leporidae | *Sylvilagus bachmani* | Brush rabbit |
| | Lagomorpha | Leporidae | *Sylvilagus floridanus* | Eastern cottontail |
| | Rodentia | Aplodontiidae | *Aplodontia rufa* | Mountain beaver |
| | Rodentia | Cricetidae | *Arborimus longicaudus* | Red tree vole |
| | Rodentia | Cricetidae | *Arborimus pomo* | Sonoma tree vole |
| | Rodentia | Cricetidae | *Microtus californicus* | California vole |
| | Rodentia | Cricetidae | *Microtus chrotorrhinus* | Rock vole |
| | Rodentia | Cricetidae | *Microtus longicaudus* | Long-tailed vole |
| | Rodentia | Cricetidae | *Microtus montanus* | Montane vole |
| | Rodentia | Cricetidae | *Microtus ochrogaster* | Prairie vole |
| | Rodentia | Cricetidae | *Microtus richardsoni* | North American water vole |
| | Rodentia | Cricetidae | *Myodes californicus* | Western red-backed vole |
| Mammalia | Rodentia | Cricetidae | *Myodes gapperi* | Southern red-backed vole |
| | Rodentia | Cricetidae | *Neotoma cinerea* | Bushy-tailed woodrat |
| | Rodentia | Cricetidae | *Neotoma fuscipes* | Dusky-footed woodrat |
| | Rodentia | Cricetidae | *Peromyscus boylii* | Brush mouse |
| | Rodentia | Cricetidae | *Peromyscus sonoriensis* | Western deer mouse |
| | Rodentia | Cricetidae | *Peromyscus truei* | Pinyon mouse |
| | Rodentia | Cricetidae | *Phenacomys intermedius* | Western heather vole |
| | Rodentia | Geomyidae | *Thomomys bottae* | Bottas pocket gopher |
| | Rodentia | Muridae | *Rattus norvegicus* | Norwegian rat |
| | Rodentia | Sciuridae | *Glaucomys sabrinus* | Northern flying squirrel |
| | Rodentia | Sciuridae | *Neotamias amoenus* | Yellow-pine chipmunk |
| | Rodentia | Sciuridae | *Neotamias siskiyou* | Siskiyou chipmunk |
| | Rodentia | Sciuridae | *Neotamias sonomae* | Sonoma chipmunk |
| | Rodentia | Sciuridae | *Neotamias speciosus* | Lodgepole chipmunk |
| | Rodentia | Sciuridae | *Neotamias townsendii* | Townsend's chipmunk |
| | Rodentia | Sciuridae | *Otospermophilus beecheyi* | California ground squirrel |

| Class | Order | Family | Scientific Name | Common Name |
|---|---|---|---|---|
| Mammalia | Rodentia | Sciuridae | *Sciurus griseus* | Western grey squirrel |
| | Rodentia | Sciuridae | *Tamiasciurus douglasii* | Douglas squirrel |
| | Rodentia | Sciuridae | *Tamiasciurus hudsonicus* | American red squirrel |
| Reptilia | Squamata | Anguidae | *Elgaria coerulea* | Northern alligator lizard |
| | Squamata | Anguidae | *Elgaria multicarinata* | Southern alligator lizard |
| | Squamata | Colubridae | *Coluber constrictor* | North American racer |
| | Squamata | Colubridae | *Contia tenuis* | Common sharp-tailed snake |
| | Squamata | Colubridae | *Diadophis punctatus* | Ringneck snake |
| | Squamata | Colubridae | *Lampropeltis californiae* | California kingsnake |
| | Squamata | Colubridae | *Lampropeltis zonata* | California mountain kingsnake |
| | Squamata | Colubridae | *Masticophis lateralis* | Striped racer |
| | Squamata | Colubridae | *Masticophis taeniatus* | Striped whipsnake |
| | Squamata | Colubridae | *Pituophis catenifer* | Gophersnake |
| | Squamata | Colubridae | *Thamnophis couchii* | Couch's garter snake |
| | Squamata | Colubridae | *Thamnophis elegans* | Terrestrial garter snake |
| | Squamata | Colubridae | *Thamnophis sirtalis* | Common garter snake |
| | Squamata | Phrynosomatidae | *Sceloporous occidentalis* | Western fence lizard |
| | Squamata | Viperidae | *Crotalus oreganus* | Western rattlesnake |
| | Squamata | Xantusiidae | *Xantusia sierrae* | Sierra night lizard |
| | Testudines | Emydidae | *Chrysemys picta* | Painted turtle |
| Actinopterygii | Salmoniformes | Salmonidae | *Oncorhynchus kisutch* | Coho salmon |
| | Clupeiformes | Clupeidae | *Clupea pallasii* | Pacific herring |
| Arachnida | Araneae | Antrodiaetidae | *Antrodiaetus pacificus* | Mygalomorph spider |
| | Araneae | Cheiracanthiidae | *Cheiracanthium mildei* | Northern yellow sac spider |
| | Araneae | Dictynidae | *Dictyna major* | Mesh web weaver |
| | Araneae | Dysderidae | *Dysdera crocata* | Woodlouse spider |
| | Araneae | Linyphiidae | *Bathyphantes gracilis* | Grassland spider |
| | Araneae | Linyphiidae | *Diplostyla concolor* | Dwarf spider |
| | Araneae | Linyphiidae | *Frontinella communis* | Bowl-and-doily spider |
| | Araneae | Linyphiidae | *Lepthyphantes leprosus* | Sheetweb spider |
| | Araneae | Pholcidae | *Pholcus phalangioides* | Cellar spider |

| Class | Order | Family | Scientific Name | Common Name |
| --- | --- | --- | --- | --- |
| Arachnida | Araneae | Salticidae | *Evarcha proszynskii* | Jumping spider |
| | Araneae | Salticidae | *Salticus scenicus* | Zebra jumping spider |
| | Araneae | Theridiidae | *Latrodectus hesperus* | Western black widow |
| | Araneae | Theridiidae | *Parasteatoda tepidariorum* | Common house spider |
| | Araneae | Theridiidae | *Steatoda triangulosa* | Triangulate cobweb spider |
| | Araneae | Ulobridae | *Hyptiotes gertschi* | Gertschs triangleweaver |
| | Ixodida | Ixodidae | *Ixodes scapularis* | Deer tick |
| Chilopoda | Lithobiomorpha | Lithobiidae | *Lithobius forficatus* | Brown centipede |
| | Scolopendromorpha | Scolopocryptopidae | *Scolopocryptops spinicaudus* | Centipede |
| Clitellata | Opisthopora | Lumbricidae | *Aporrectodea caliginosa* | Grey worm |
| | Opisthopora | Lumbricidae | *Lumbricus rubellus* | Red worm |
| | Opisthopora | Lumbricidae | *Lumbricus terrestris* | Common earthworm |
| Diplopoda | Julida | Julidae | *Cylindroiulus caeruleocinctus* | Millipede |
| | Julida | Julidae | *Ophyiulus pilosus* | Furry snake millipede |
| Gastropoda | Architaenioglossa | Viviparidae | *Cipangopaludina chinensis* | Chinese mystery snail |
| | Gastropod | Agriolimacidae | *Deroceras laeve* | Marsh slug |
| | Stylommatophora | Ariolimacidae | *Ariolimax columbianus* | Pacific banana slug |
| | Stylommatophora | Arionidae | *Arion ater* | Large black slug |
| | Stylommatophora | Xanthonychidae | *Monadenia fidelis* | Pacific sideband snail |
| Insecta | Blattodea | Blattidae | *Periplaneta americana* | American cockroach |
| | Coleoptera | Coccinellidae | *Harmonia axyridis* | Asian ladybeetle |
| | Hemiptera | Cicadidae | *Okanagana rimosa* | Says cicada |
| | Lepidoptera | Erebidae | *Hyphantria cunea* | Fall webworm moth |
| | Lepidoptera | Erebidae | *Orgyia antiqua* | Rusty tussock moth |
| | Lepidoptera | Noctuidae | *Mythimna unipuncta* | Armyworm moth |
| | Lepidoptera | Noctuidae | *Noctua pronuba* | Large yellow underwing moth |
| | Lepidoptera | Noctuidae | *Spodoptera frugiperda* | Fall armyworm |
| | Lepidoptera | Noctuidae | *Spodoptera ornithogalli* | Yellow-striped armyworm |
| | Lepidoptera | Noctuidae | *Trichoplusia ni* | Cabbage looper |
| | Lepidoptera | Noctuidae | *Xestia c-nigrum* | Setaceous Hebrew character moth |

| Class | Order | Family | Scientific Name | Common Name |
|---|---|---|---|---|
| Insecta | Lepidoptera | Saturniidae | *Hyalophora cecropia* | Cecropia moth |
| | Lepidoptera | Sphingidae | *Manduca sexta* | Tobacco hornworm |
| | Odonata | Libellulidae | *Plathemis lydia* | Common whitetail |
| | Orthoptera | Acrididae | *Chorthippus curtipennis* | Meadow grasshopper |
| | Orthoptera | Acrididae | *Dissosteira carolina* | Carolina grasshopper |
| | Orthoptera | Acrididae | *Melanoplus bivittatus* | Two-striped grasshopper |
| | Orthoptera | Acrididae | *Melanoplus scudderi* | Scudder's short-winged grasshopper |
| | Orthoptera | Acrididae | *Schistocerca shoshone* | Green bird grasshopper |
| | Orthoptera | Stenopelmatidae | *Stenopelmatus fuscus* | Jerusalem cricket |
| | Orthoptera | Tettigoniidae | *Scudderia furcata* | Fork-tailed bush katydid |
| Malacostraca | Decopoda | Astacidae | *Pacifastacus leniusculus* | Signal crayfish |
